# Phandango: an interactive viewer for bacterial population genomics

**DOI:** 10.1101/119545

**Authors:** James Hadfield, Nicholas J. Croucher, Richard J Goater, Khalil Abudahab, David M Aanensen, Simon R. Harris

## Abstract

**Summary:** Fully exploiting the wealth of data in current bacterial population genomics datasets requires synthesising and integrating different types of analysis across millions of base pairs in hundreds or thousands of isolates. Current approaches often use static representations of phylogenetic, epidemiological, statistical and evolutionary analysis results that are difficult to relate to one another. Phandango is an interactive application running in a web browser allowing fast exploration of large-scale population genomics datasets combining the output from multiple genomic analysis methods in an intuitive and interactive manner.

**Availability:** Phandango is a web application freely available for use at https://jameshadfield.github.io/phandango and includes a diverse collection of datasets as examples. Source code together with a detailed wiki page is available on GitHub at https://github.com/jameshadfield/phandango

## 1 INTRODUCTION

Bacterial population genomics has advanced rapidly in terms of numbers of genomes sequenced, with recent publications involving analyses of hundreds or even thousands of bacterial genomes. Such studies often base their understanding upon a phylogenetic tree, onto which epidemiological, comparative genomic and phenotypic data can be mapped. In bacterial species which undergo homologous recombination, horizontal sequence transfer means that whole-genome phylogenies often have to be adjusted to mitigate the confounding effects of recombination using methods such as Gubbins (Croucher *et al*., 2015) or BRAT NextGen (Marttinen *et al*., 2011). These methods also predict regions of horizontally imported DNA in the genome of each bacterial isolate, which can only be practically interpreted when displayed in the context of the phylogeny. An alternative approach to large-scale comparative genomics is to investigate the distribution of the pan-genome across a set of isolates using software such as ROARY (Page *et al*., 2015). Finally, increasing sample sizes have opened the way for genetic and phenotypic data to be combined in genome-wide association studies (GWAS) using programs such as PLINK or SEER (Lees *et al*., 2016; Purcell *et al*., 2007). These approaches have proved successful in identifying serotype switching within populations or finding variants associated within antimicrobial resistance (Croucher *et al*., 2011; Chewapreecha *et al*., 2014).

Increasingly, web application development provides us with methods to link and visualise complex genomic data interactively (Argimón *et al*., 2016) However, recombination, pan-genome and GWAS analyses all produce large amounts of output data that are typically explored separately in visually distinct styles, relative to a phylogeny, a reference sequence, or both. Currently exploratory analyses are often represented as single static images that provide a simple overview but do not allow visual investigation of the data or the ability to relate output from multiple analyses to one another. The ability to interactively visualize such complex and information rich datasets would allow clearer interpretation and facilitate novel biological discoveries.

Phandango is an interactive web application which runs directly in web browsers. Data are uploaded by dragging and dropping files onto the browser window and analysed client-side such that no data is transferred to servers. Uploaded data can by added to or replaced by dropping new files at any time. Figure 1A illustrates the resulting grid layout produced when a phylogenetic tree, an associated metadata file, a reference sequence annotation file and the output from Gubbins and BRATNextGen are simultaneously uploaded into Phandango. The grid is defined by a phylogeny and a reference sequence annotation; output of other analyses are then drawn within this framework. The resulting visualization is fully interactive, allowing users to manipulate and zoom both the phylogeny and along the length of the reference sequence using intuitive controls. The space allocated to panels within the grid can be easily adjusted by dragging. The framework allows loci of interest highlighted by any of the supported population genomic analysis data formats to be easily cross-referenced with functional information associated with the reference genome. This means that multiple population genomic analyses can be interactively compared in a single environment. Phandango is written in modern JavaScript and uses the PhyloCanvas library (https://github.com/PhyloCanvas/PhyloCanvas) for interactive display of phylogenies.

## 2. DATA TYPES ACCEPTED

Phandango is versatile in the types of data format which can be displayed. For a detailed explanation of the supported data formats please see https://github.com/jameshadfield/phandango/wiki/Input-data-formats. Briefly, phylogenies are expected in Newick format, recombination, GWAS and pan genome data are expected in the default output formats of the software that produced them (currently supported software are Gubbins, BRATNextGen, PLINK, SEER and ROARY), genome annotations are expected in GFF3 format and metadata in simple CSV format. Since all of these inputs are simple text files, it is relatively simple for any custom data structure may to be converted by the user into one of these formats and subsequently displayed.

**Fig. 1.**
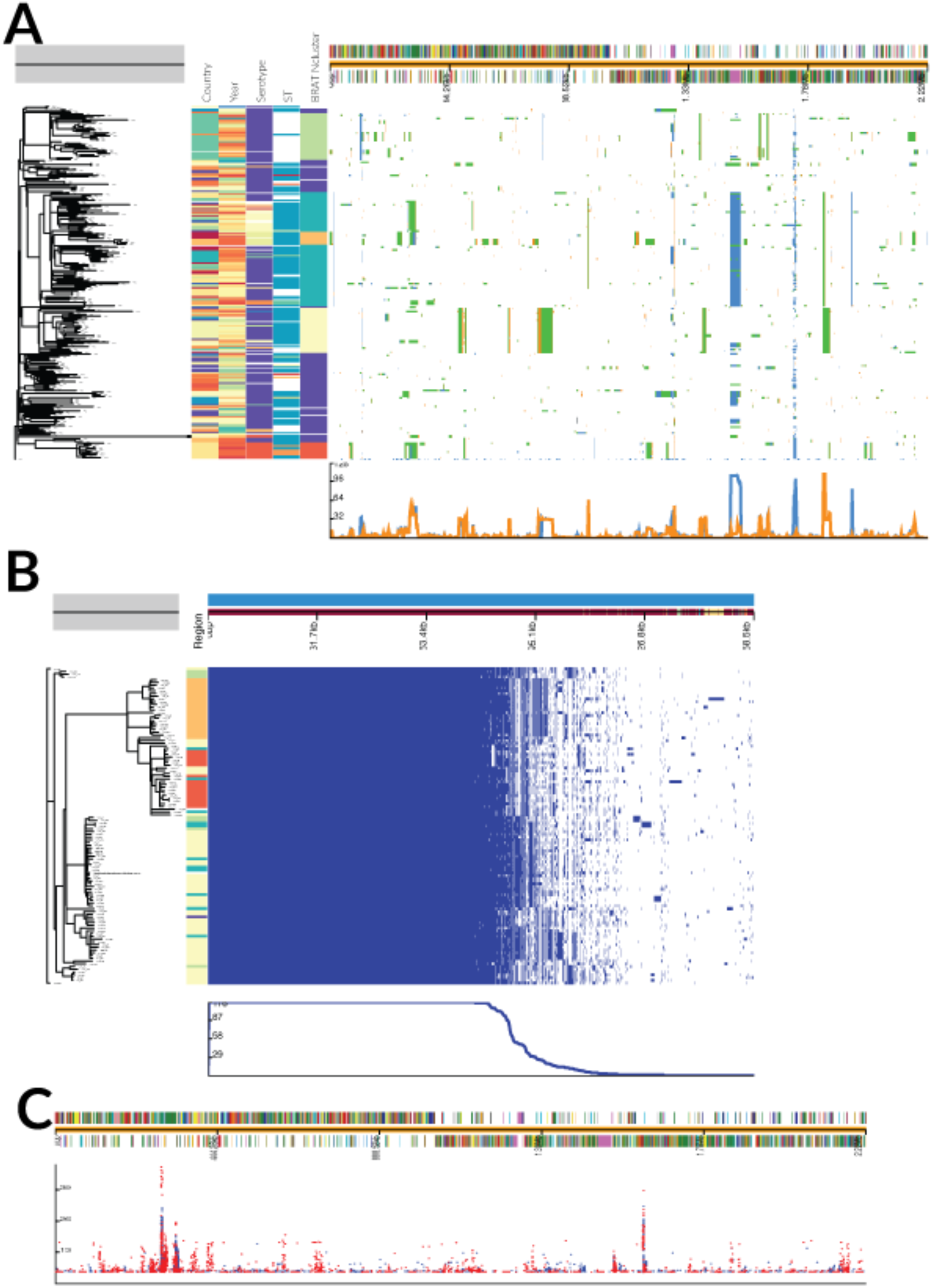
Screenshots of three different datasets of *Streptococcus pneumoniae* genomes (all available as online examples). (A) Visualisation of recombinations inferred using an *S. pneumoniae* PMEN1 dataset using Gubbins (blue blocks) & BRATNextGen (yellow blocks); green blocks represent overlap between methods. (B) ROARY pan-genome analysis. (C) GWAS results generated by PLINK relative to the genome of *S. pneumoniae* ATCC 700669.

## 3. USER INTERFACE

Phandango initially presents the entirety of the user’s data (normally consisting of the entire phylogeny and the entire reference sequence or pan genome) simultaneously, as in Fig. 1. The exact nature of the layout depends on the data loaded – for instance, one can view simply a phylogeny and associated metadata, or a genome annotation together with GWAS results without a phylogeny (Fig. 1c). The user can then quickly and easily zoom into regions of the genomic data, effectively expanding the view horizontally to focus on particular genomic loci. This allows rapid biological interpretation of complex data by quickly viewing the genomic regions of interest in greater detail. Combined with the ability to interact with the phylogeny by zooming to focus on particular leaf nodes or selecting and drawing sub-trees, the user can, for example, explore lineage specific recombination or pan-genome profiles and compare these results against the overall dataset. Hovering over the genome annotation (top) or the metadata (between the phylogeny and the genomic information) displays any annotation associated with that data. A line graph is automatically generated and displayed under the genomic information panel. Depending on the data type displayed, the line graph represents either the recombination prevalence along the sequence, or the number of isolates containing a particular gene. If sub clades are selected on the tree, a second line graph is overlaid showing the same data for the selected taxa. In this way features of sub lineages may be easily compared to those of the overall dataset. If both Gubbins and BRAT NextGen data are loaded then the user may toggle between the results of the two methods separately or a merged output showing their overlap (Figure 1A).

Further options, including the ability to output the view as a vector image, are available online at https://github.com/jameshadfield/phandango/wiki.

## 4. CONCLUSIONS

Phandango is an intuitive, user-friendly application which requires no installation or command line knowledge. It allows rapid viewing and interactive exploration of large genomic datasets and aids biological understanding of complex data through linking the output of multiple genomic analysis methods into a single, intuitive interface.

## REFERENCES

Argimón, S. et al. (2016) Microreact: visualizing and sharing data for genomic epidemiology and phylogeography. Microbial Genomics, 10, 199.

Chewapreecha, C. et al. (2014) Comprehensive Identification of Single Nucleotide Polymorphisms Associated with Beta-lactam Resistance within Pneumococcal Mosaic Genes. PLoS Genet, 10, e1004547–11.

Croucher, N.J. et al. (2015) Rapid phylogenetic analysis of large samples of recombinant bacterial whole genome sequences using Gubbins. Nucleic Acids Research, 43, e15–e15.

Croucher, N.J. et al. (2011) Rapid Pneumococcal Evolution in Response to Clinical Interventions. Science, 331, 430–434.

Lees, J.A. et al. (2016) Sequence element enrichment analysis to determine the genetic basis of bacterial phenotypes.

Marttinen, P. et al. (2011) Detection of recombination events in bacterial genomes from large population samples. Nucleic Acids Research, 40, e6–e6.

Page, A.J. et al. (2015) Roary: rapid large-scale prokaryote pan genome analysis. Bioinformatics, 31, 3691–3693.

Purcell, S. et al. (2007) PLINK: A Tool Set for Whole-Genome Association and Population-Based Linkage Analyses. Am. J. Hum. Genet., 81, 559–575.

